# Infectious potential and circulation of SARS-CoV-2 in wild rats

**DOI:** 10.1101/2024.12.19.629569

**Authors:** Kevyn Beissat, Virginie Lattard, Evelyne Picard-Meyer, Ambre Fafournoux, Sionfoungo Daouda Soro, Alexandre Servat, Françoise Vincent-Hubert, Franck Boué, Nolan Chatron, Elodie Monchâtre-Leroy, Marine Wasniewski

**Author notes:** Corresponding authors: (KB), (MW).

## Abstract

Since the beginning of the severe acute respiratory syndrome coronavirus 2 (SARS-CoV-2) pandemic, a wide range of animal species (pets, mink…) have been naturally infected with this betacoronavirus. The emergence of new variants has increased the ability of SARS-CoV-2 to infect species that were not susceptible to the “original” SARS-CoV-2, such as mice and rats.

This work attempted to evaluate the role of urban rats in the SARS-CoV-2 transmission by combining surveillance studies of rat populations in urban environments, *in vivo* experimental inoculation of SARS-CoV-2 and comparative viral-receptor interaction in *silico* analyses.

We studied the circulation of SARS-CoV-2 in wild *Rattus norvegicus* (n=401) captured in urban areas and sewage systems of several French cities. Except for 3 inconclusive samples (2/75 from Bordeaux and 1/261 from Lyon) none of the 353 sera tested showed anti-SARS-CoV-2 antibodies by microsphere immunoassay. However, the 3 inconclusive sera samples were negative by virus neutralisation assay. No SARS-CoV-2 viral RNA was detected in all lungs collected from the 401 captured urban brown rats. In complement, four rat groups (two wild-type colonies, *Rattus norvegicus* and *Rattus rattus*, and two laboratory strains, Sprague-Dawley and Wistar) were inoculated with the SARS-CoV-2 Omicron BA.5. At 4 days post-inoculation, no infectious viral particles were detected in the lungs and upper respiratory tract (URT) while viral RNA was detected at a low level only in the URT of all groups. In addition, seroconversion was observed 14 days after inoculation in the four groups. By molecular modelling, the Omicron BA.5 receptor binding domain (RBD) had lower affinities for *Rattus norvegicus* and *Rattus rattus* ACE2 than *Homo sapiens* ACE2.

Based on these results the SARS-CoV-2 Omicron BA.5 was unable to infect laboratory and wild type rats. In addition, *Rattus norvegicus* collected for this study in different areas of France were not infected with SARS-CoV-2.

## Introduction

In late December 2019, a novel coronavirus called severe acute respiratory coronavirus 2 (SARS-CoV-2) was identified as the causative agent of COVID-19 disease in Wuhan, China [1]. This virus, which belongs to the betacoronavirus genus, rapidly spread in the human population and caused numerous deaths with serious repercussions on global health. Shortly after the onset of the pandemic, the first cases of natural animal infections occurred [2]. Indeed, several animal species, including pets (dogs and cats), farm animals (minks), zoo animals (tigers and lions), and wild animals (otters and white-tailed deer) were reported to be naturally infected with this coronavirus [3–8]. This phenomenon warrants careful consideration as it may increase the risk of viral mutation and subsequent transmission of potentially more pathogenic viruses to humans. This phenomenon has already been observed in mink and white-tailed deer species [9,10]. Mice and rats were not susceptible to infection with the ‘’original” SARS-CoV-2 [11–13]. However, the emergence of new variants of concern (VOCs) such as Alpha (B.1.1.7), Beta (B.1.351) and Omicron (B.1.1.529, BA.5…) variants with favourable mutations in the spike protein seems to modify the viral host range of this virus. Indeed, infectious viral particles have been detected in the nasal turbinates and lungs of C57BL/6J mice and Sprague-Dawley rats infected with the Alpha, Beta or Delta (B.1.617.2) VOC [12,14,15]. In addition, it was demonstrated that the Delta variant is unable to infect wild mice, whereas the Beta variant can spread through close contact between infected and naive BALB/c mice [16] and between Sprague-Dawley rats [14]. These results suggest that mutations in the spike protein such as the N501Y mutation may enhance the ability of the virus to infect new species, including rodents.

The SARS-CoV-2 genome has been widely found in wastewater [17], an environment, where wild rats roam freely. These findings raise concerns about the potential transmission of the virus to urban rodents. While the presence of infectious SARS-CoV-2 particles in wastewater has not yet been demonstrated, a recent experimental study, highlighted the survival of SARS-CoV-2 infectious particles in unfiltered/filtered raw sewage, and secondary effluent for several hours (between 10 and 18 hours) [18]. The emergence of new variants and the circulation of SARS-CoV-2 viral RNA in sewage level up the risk of infection of urban rats. Therefore, the possibility of urban rodents becoming a new animal reservoir for the virus, with potential spillover to humans, should not be overlooked. In addition, a study conducted on wild rats captured in urban areas of the USA, close to wastewater, detected partial SARS-CoV-2 genome in the lungs of *Rattus norvegicus* [15]. The presence of antibodies against SARS-CoV-2 has also been demonstrated in brown rats (*Rattus norvegicus*) in the USA, the United-Kingdom and Canada [15,19,20].

In this context, this work attempted to evaluate the role of wild rats in the SARS-CoV-2 transmission by combining surveillance studies of rat populations in urban environments across various French cities, *in vivo* experimental inoculation of SARS-CoV-2 into both wild and laboratory rats and comparative viral-receptor interaction in *silico* analyses.

## Results

### Field monitoring of SARS-CoV-2 infections in urban rat populations

A total of 401 wild rats were collected from six French cities (Fig 1 and Table 1), distributed homogenously across the country, with human populations ranging between 200,000 to 1,500,000 inhabitants and average population densities between 2,000 and 10,000 inhabitants per square kilometre: Besancon (4.7% of the rats), Bordeaux (19.2%), Lyon (67.1%), Marseille (3.0%), Nancy (5.0%) and Nantes (1.0%) between January 2022 and July 2023 (Fig 2). These rats were trapped in sewers (29.2%), in social housing courtyards with direct connection to the sewer network (58.3%) or in urban parks without connection to the sewer network (12.5%). Within the sample, 57.2% were males and the median weight of trapped rats was 266.7g (95% CI: 243.8-287.9g), with values ranging from 26g to 600g (excluding Marseille rats, which could not be weighed) (Table 1). All the 401 wild rats have been confirmed to belong to the species *Rattus norvegicus* through morphometric measurements, as well as by amplification and sequencing of the cytochrome b gene in cases of uncertainty.

**Fig 1.**
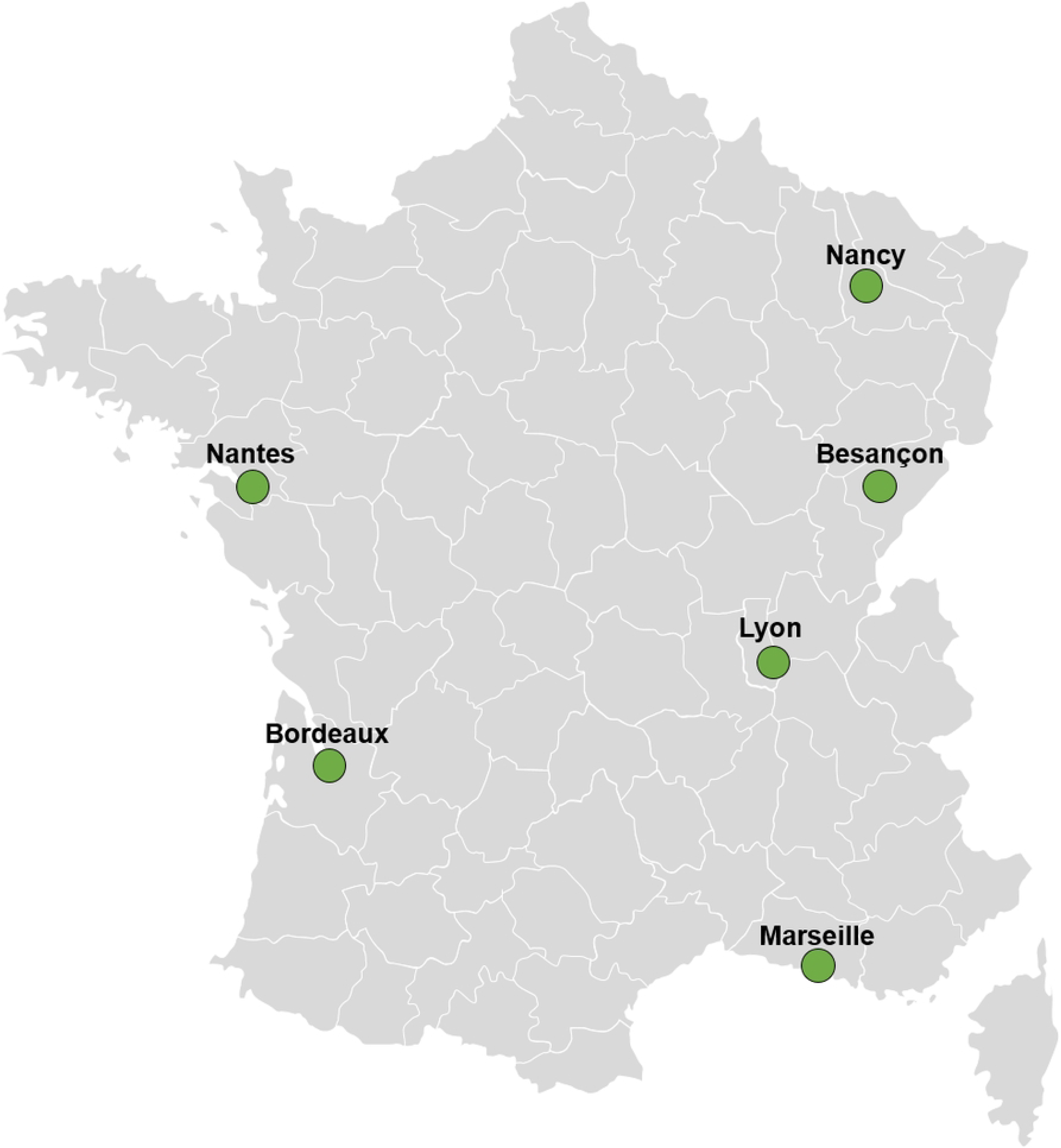
Localisation of the different urban areas of capture in France.

**Fig 2.**
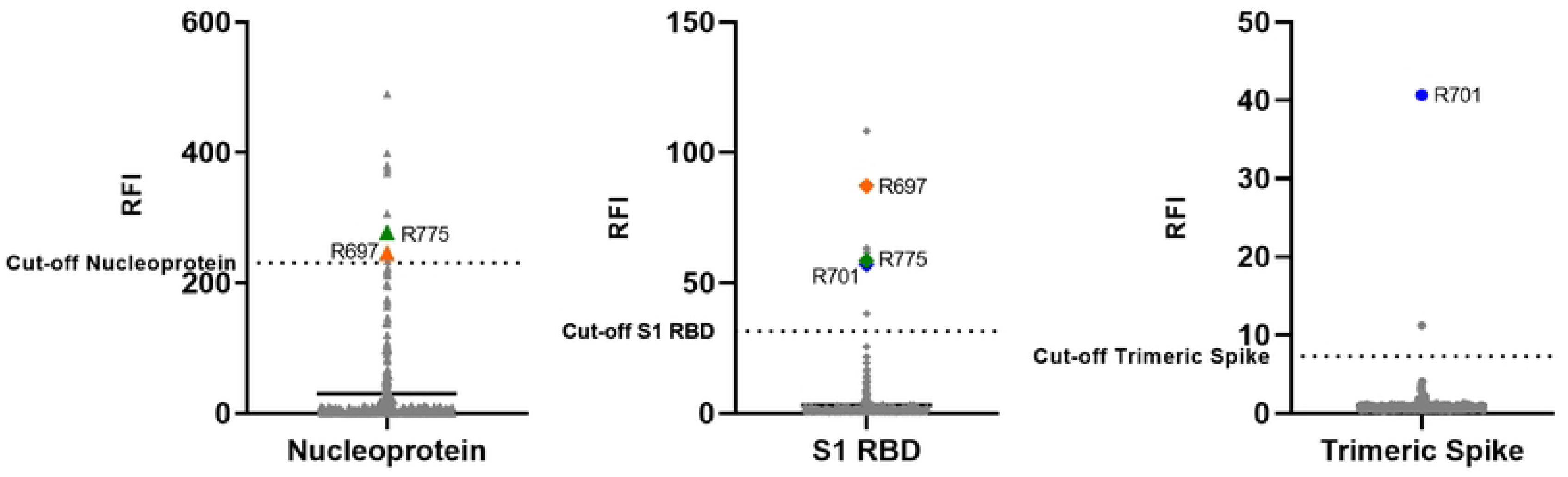
Detection of sera anti-SARS-CoV-2 antibodies in the urban rats using microsphere immunoassay (MIA). Antibodies were detected using three SARS-CoV-2 antigens (the nucleoprotein, the subunit 1 (S1) receptor binding domain (RBD) of the spike and the trimeric spike). Antibody levels were expressed as Relative Fluorescence Intensities (RFI) to control antigen. The negative group (the whole population) was used to determine the cut-off (mean + 3 standard deviation). In all graphs, mean-values are presented. Seroconversion was defined as the detection of antibodies against the three antigens, while results were considered inconclusive if antibodies were detected against two of the three antigens.

**Table 1.**
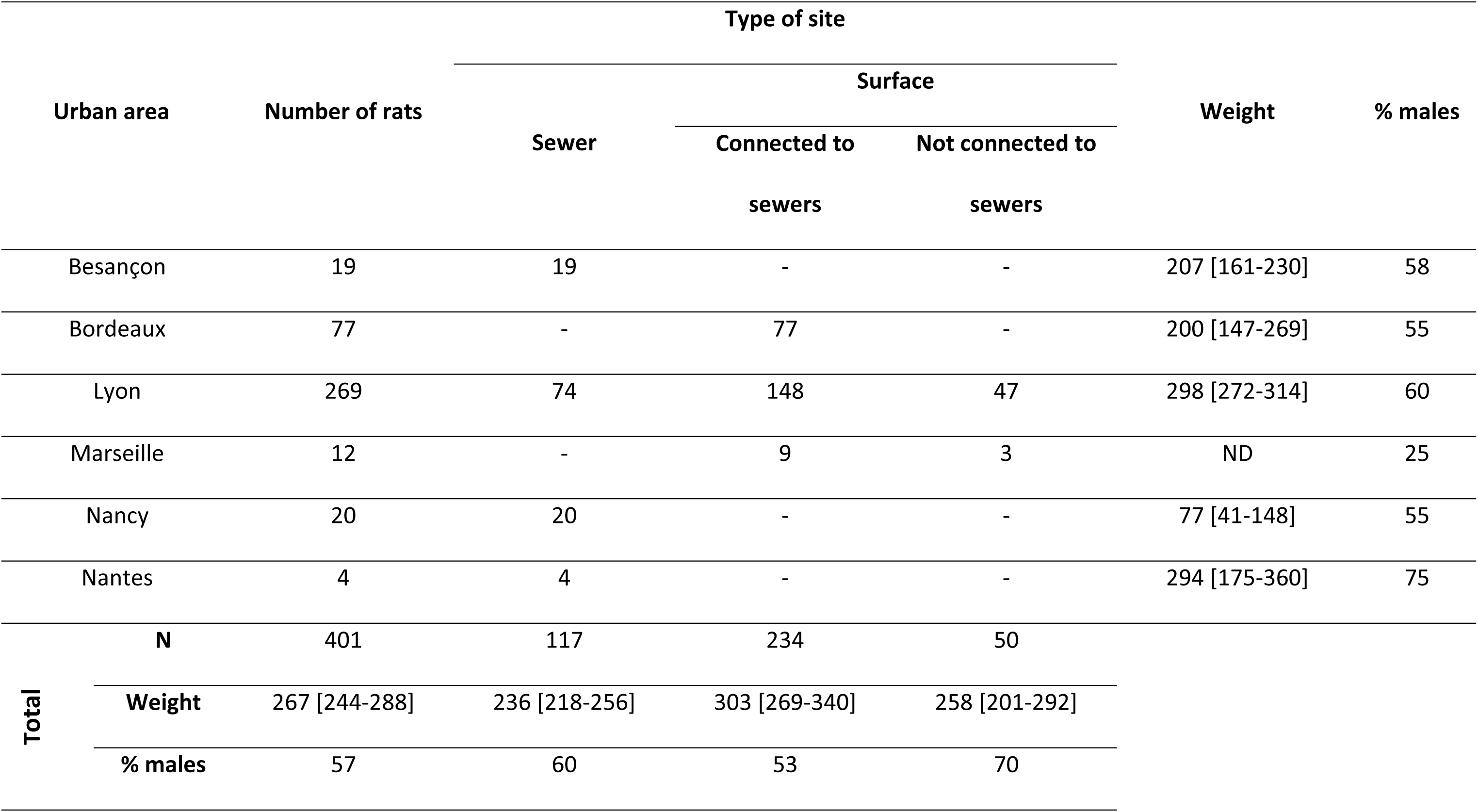
Characterization of wild urban rat populations captured in these cities as part of the surveillance study on SARS-CoV-2 infection in rats.

**Table 2.** Comparison of the results obtained with the microsphere immunoassay and the seroneutralisation methods for the four groups of rats.

| Group | Method | Rat identifier |  |  |  |  |  |  |  |
| --- | --- | --- | --- | --- | --- | --- | --- | --- | --- |
|  |  | R4 | R5 | R6 | R7 | R8 | R9 | R10 | R11 |
| *Rattus norvegicus* (wild) | MIA | Yes | Inc | Yes | Yes | Inc | Inc | Yes | Inc |
|  | SN | Yes | Yes | Yes | No | No | No | No | Yes |
|  |  | R11 | R12 | R13 | R14 | R15 | R16 | R20 | - |
| *Rattus rattus* | MIA | Inc | Inc | Yes | Inc | Inc | Yes | Yes | - |
|  | SN | No | No | Yes | No | No | No | No | - |
|  |  | R14 | R15 | R16 | R17 | R18 | R19 | R20 | - |
| Sprague-Dawley | MIA | Yes | Yes | Yes | Yes | Yes | Yes | Yes | - |
|  | SN | No | Yes | No | No | Yes | Yes | Yes | - |
|  |  | R14 | R15 | R16 | R17 | R18 | R19 | R20 | - |
| Wistar | MIA | Yes | Yes | Yes | Yes | Yes | Yes | Yes | - |
|  | SN | Yes | No | Yes | Yes | Yes | No | Yes | - |

MIA: Microsphere immunoassay and SN: seroneutralisation. “Yes”: presence of a seroconversion, “No”: absence of a seroconversion and “Inc”: inconclusive results (for MIA results presented in Fig 5).

401 lungs and 353 sera (75 from Bordeaux, 261 from Lyon, 12 from Marseille and 5 from Nancy) were collected from the 401 rats captured. All animals were found to be RT-PCR negative for SARS-CoV-2 in the lungs, while wastewaters collected in sewers were positive depending on the city (Besançon Ct = 28 – 37; Lyon = 33-35; Nantes= 28 – 36; Nancy = 34 – 40). Presence of anti-SARS-CoV-2 antibodies were thus screened in 353 sera (for some rats, blood was not obtained) by MIA. To determine the cut-off of positivity, the entire rat population was used, considering the low percentage of natural infection of *Rattus norvegicus* and the absence of pre-pandemic sera [21]. None of the 353 tested sera were positive for the three SARS-CoV-2 antigens and inconclusive results were obtained for 3 samples (Fig 2). Two of these samples (R701 and R775) were collected in Bordeaux, from rats trapped in a social housing courtyard with direct connection to the sewer network, and the third sample (R697) was collected in Lyon, on a rat trapped in a sewer. These 3 samples were collected in June 2023 when the XBB 1.5 variant circulated in French human population [22]. Samples having inconclusive result were tested by seroneutralisation (SN) assay using the SARS-CoV-2 D614G as the challenge virus. All samples were found negative.

### Experimental *in vivo* inoculation of SARS-CoV-2 in laboratory and wild rats

Concurrently, we performed *in vivo* experiments to evaluate the susceptibility of both laboratory and wild rats to Omicron BA.5 variant inoculation. Since no significant differences in response were observed between the three inoculated wild *Rattus norvegicus* colonies (A, B and C), data from these colonies were combined for simplified graphical representation.

#### Clinical signs

No clinical signs or significant variations in weight (p>0.05) were observed between the negative groups and the Omicron BA.5-inoculated groups at 4 and 14 dpi whether in laboratory rats or wild *Rattus norvegicus* or *Rattus rattus* (Fig 3).

**Fig 3.**
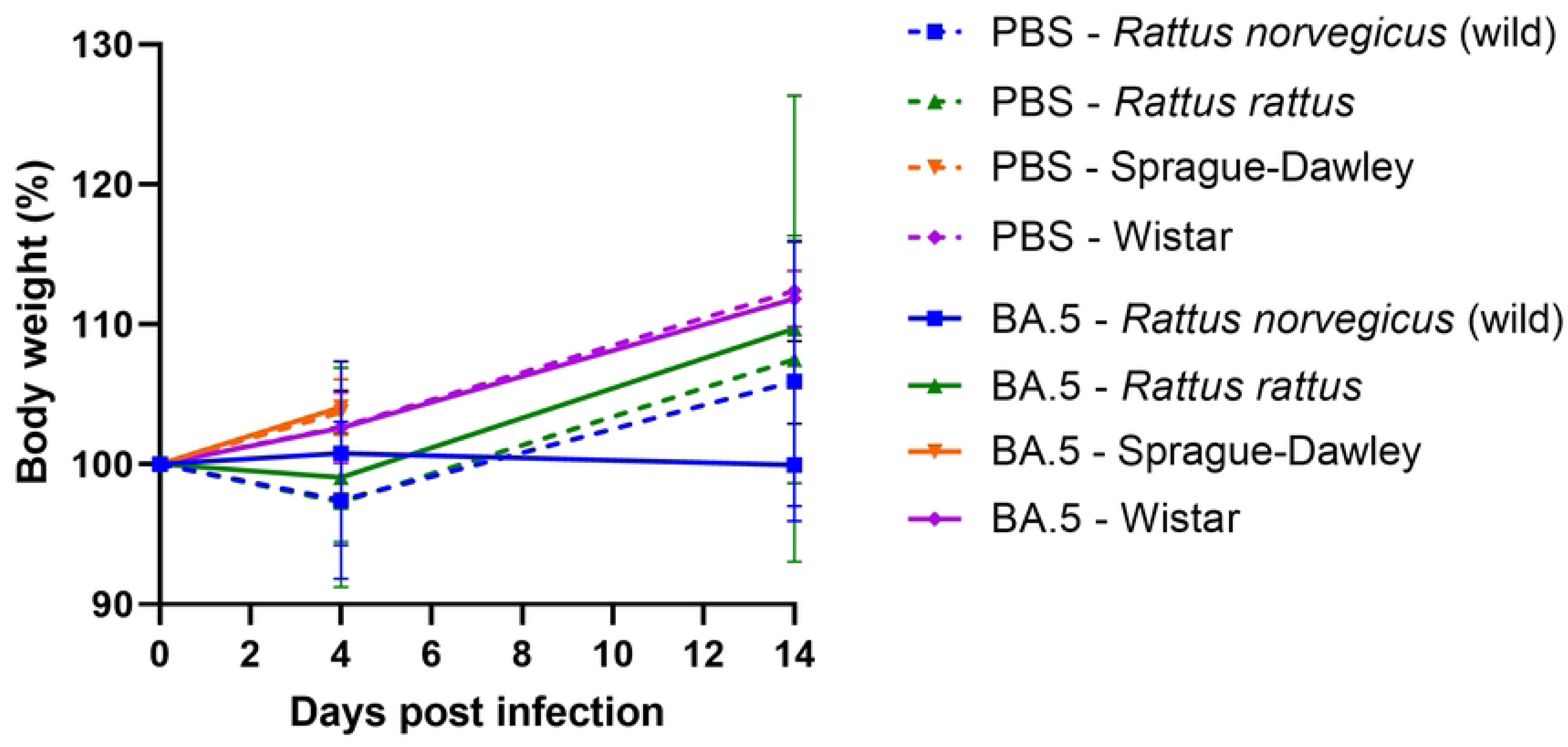
Body weight evolution of the PBS inoculated rats *versus* the Omicron BA.5 inoculated rats for the four groups. The means are represented ± one standard deviation. Data for Sprague-Dawley at D14, are missing. PBS: groups control.

#### Detection of viral RNA, infectious viral particles and antibodies anti-SARS-CoV-2 in inoculated rats

Viral RNA was detected at 4 dpi in the upper respiratory tract (URT) of all groups, including both wild and laboratory rats (Fig 4). The mean viral load for Sprague-Dawley and Wistar rats was approximately 3.3×10^2^ and 4.1×10^2^ RNA copies/µL of URT supernatant, respectively. For wild groups, it was 8×10^2^ and 8.8×10^3^ RNA copies/µL of URT supernatant from *Rattus norvegicus* and *Rattus rattus,* respectively. No significant differences (p>0.05) were observed between any of the groups. In addition, SARS-CoV-2 RNA was not detected in the lungs of any of the rats. Furthermore, no infectious viral particles were detected in cell culture by TCID50 assays at 4 dpi in the lungs or the URT of any of the four groups of rats.

**Fig 4.**
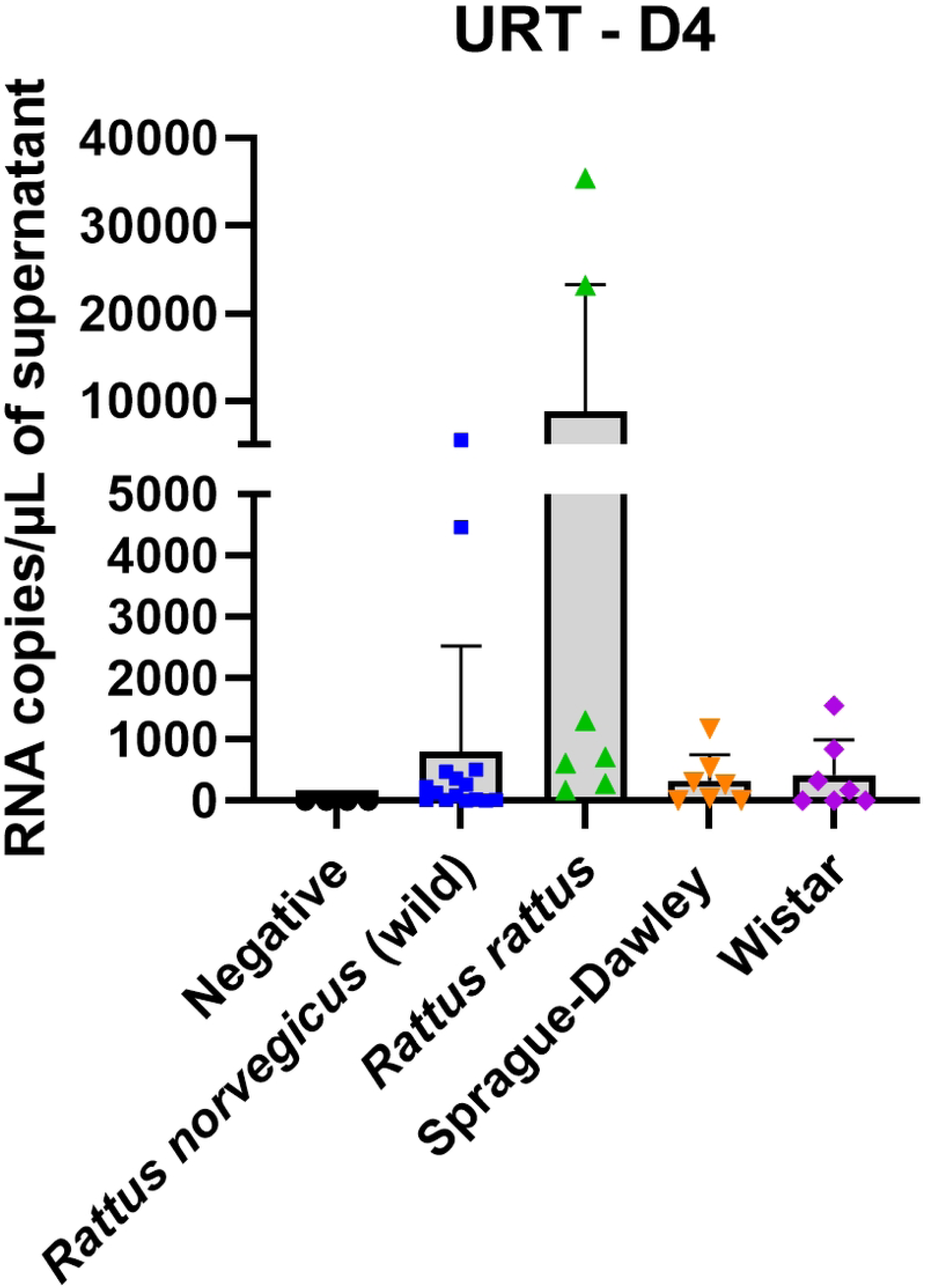
Detection of viral RNA load in the upper respiratory tract (URT) of the four rat groups at 4 days post infection (D4). The means are represented ± one standard deviation.

The presence of anti-SARS-CoV-2 antibodies in plasma was investigated by MIA at 4 and 14 dpi. At 4 dpi, no seroconversion was observed in any group of rats, whereas at 14 dpi, antibody responses were observed in the four groups of rats. However, while all Sprague-Dawley and Wistar rats seroconverted, seroconversion was only detected in 4/8 wild *Rattus norvegicus* and 4/7 *Rattus rattus* (Fig 5). To investigate the neutralisation efficiency of the detected anti-SARS-CoV-2 antibodies, a seroneutralisation (SN) assay using the Omicron BA.5 as challenge virus was performed on all plasmas (Fig 6). Four wild *Rattus norvegicus*, one *Rattus rattus* and five Sprague-Dawley and Wistar showed neutralising anti-SARS-CoV-2 antibodies up to 1/90 dilution. For both the MIA and SN methods, no antibodies were detected in the negative group. In the two laboratory rat strains and the *Rattus rattus* colony, the results showed that rats positive by SN, were also detected positive by MIA. In the wild *Rattus norvegicus* colony, seroconversion was observed in two rats (R5 and R11) by SN whereas no antibodies were detected by MIA. (Table 3). Specifically, these two samples, defined as inconclusive results, were tested positive for two out of the three SARS-CoV-2 antigens (nucleoprotein and spike trimeric antigens). In addition, all wild *Rattus norvegicus* and *Rattus rattus* sera were positive or inconclusive results.

**Fig 5.**
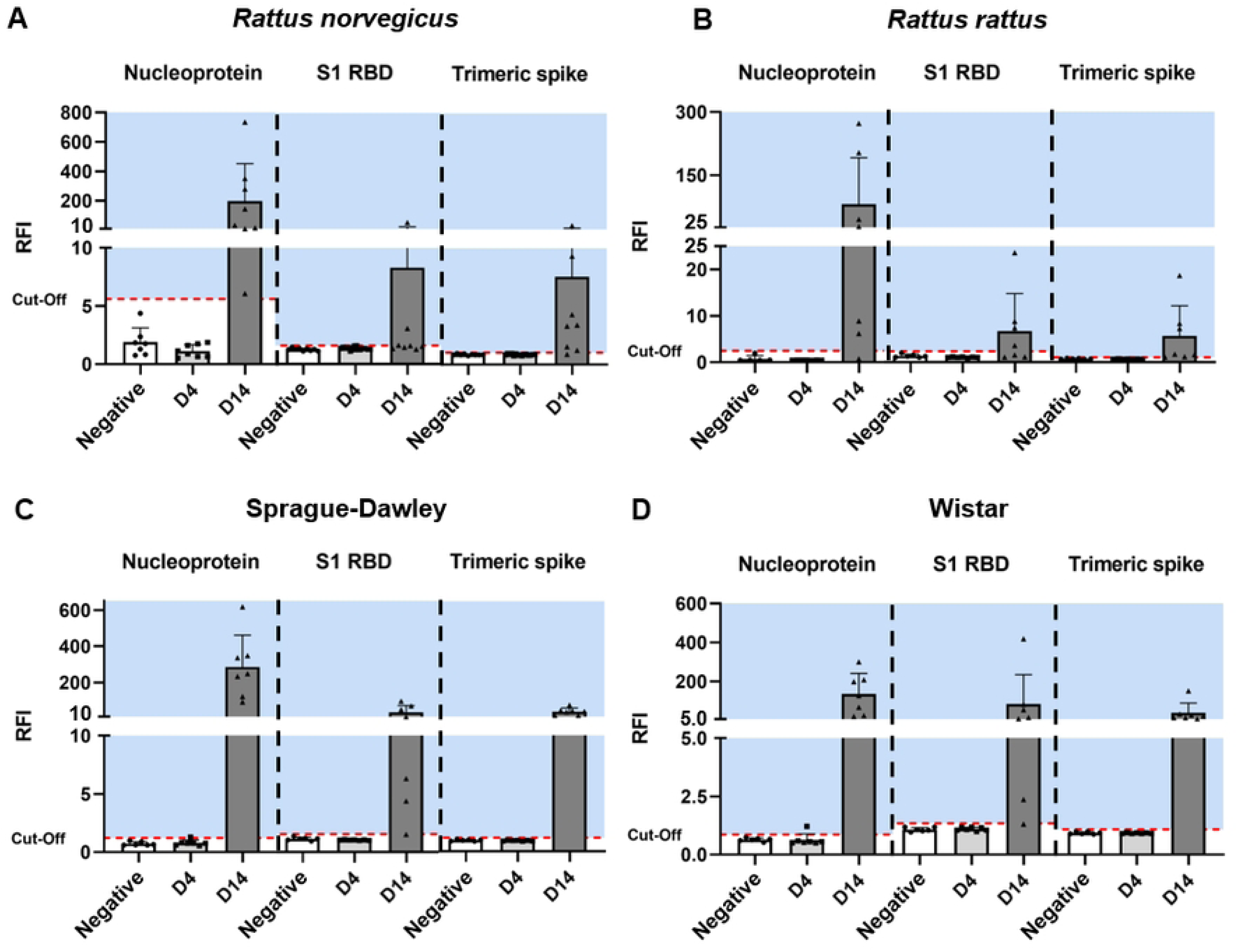
Detection of anti-SARS-CoV-2 antibodies in plasma at 4 and 14 dpi by microsphere immunoassay in the four groups of rats. Three SARS-CoV-2 antigens (the nucleoprotein, the subunit 1 (S1) receptor binding domain (RBD) of the spike and the trimeric spike) were used. The levels of anti-SARS-CoV-2 antibodies were expressed as Relative Fluorescence Intensities (RFI) to the control antigen. A: *Rattus norvegicus* (wild), B: *Rattus rattus*, C: Sprague-Dawley and D: Wistar. The negative group was used to determine the cut-off (mean + 3 standard deviation). For all graphs, the means are represented ± one standard deviation. The blue background represents the seropositivity area. Seroconversion was established when antibodies were detected with the three antigens and plasma was considered inconclusive when two of the three antigens were positive.

**Fig 6.**
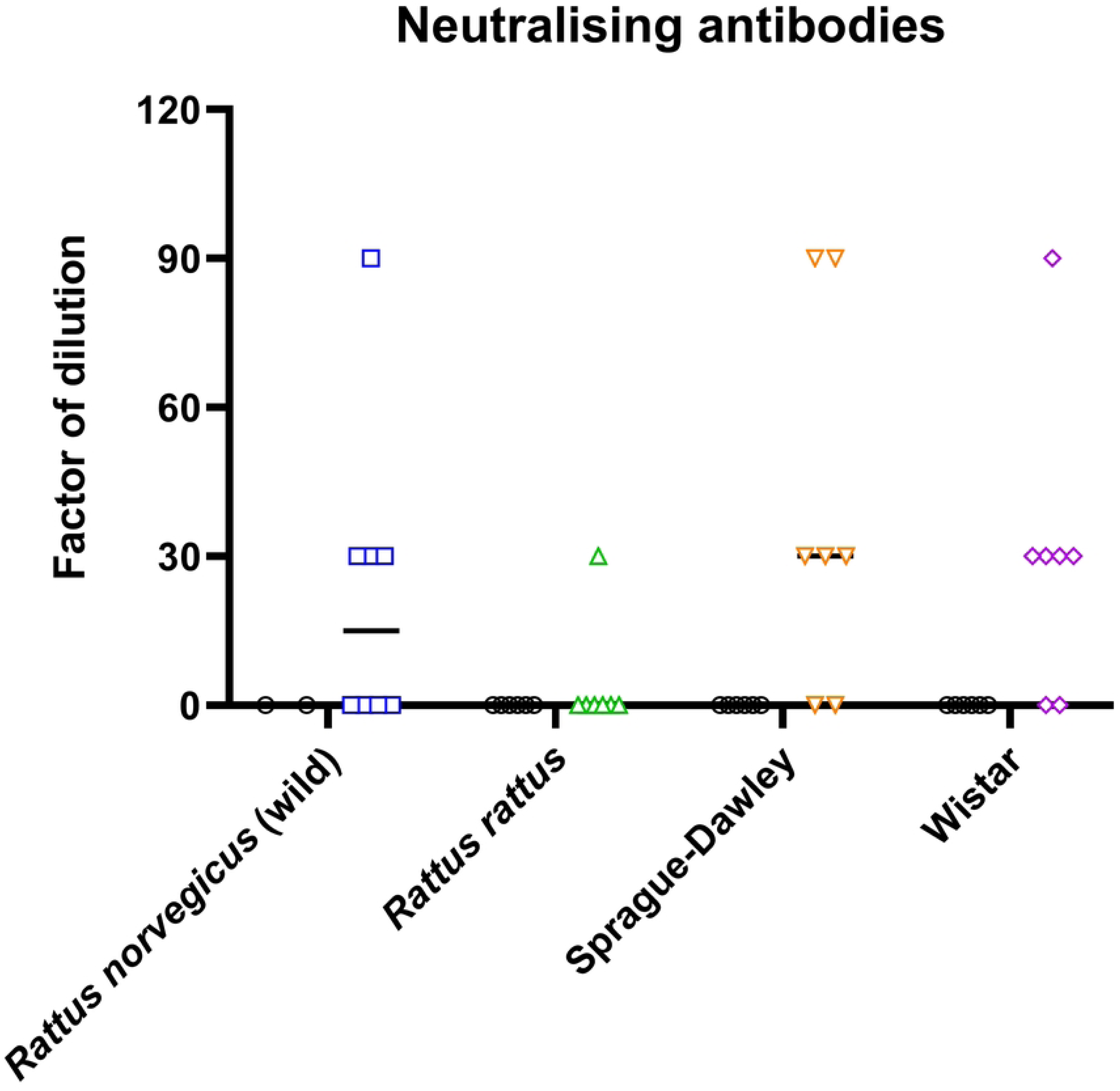
Levels of neutralising antibodies detected in the four groups of rats. *Rattus norvegicus* (wild) (blue) (n=8), *Rattus rattus* (green) (n=8), Sprague-Dawley (orange) (n=7) and Wistar (purple) (n=7) at 14 dpi. Animals from negative group are indicated by a black circle. Each point represents one rat and the mean-values are represented on the graph.

**Table 3.** Viral titers of Omicron BA.5 inoculated to the different rat groups.

| Rat group | Viral titer inoculated of Omicron BA.5 |
| --- | --- |
| *Rattus norvegicus* (wild) – group A | $10^{4.96}$ TCID<sub>50</sub>/40 μL ($10^{6.36}$TCDI<sub>50</sub>/mL) |
| *Rattus norvegicus* (wild) – group B | $10^{5.34}$ TCID<sub>50</sub>/40 μL ($10^{6.74}$TCDI<sub>50</sub>/mL) |
| *Rattus norvegicus* (wild) – group C | $10^{5.23}$ TCID<sub>50</sub>/40 μL ($10^{6.63}$TCDI<sub>50</sub>/mL) |
| *Rattus norvegicus* (wild) | $10^{5.12}$ TCID<sub>50</sub>/40 μL ($10^{6.52}$TCDI<sub>50</sub>/mL) |
| *Rattus Rattus* | $10^{4.57}$ TCID<sub>50</sub>/40 μL ($10^{5.97}$TCDI<sub>50</sub>/mL) |
| Sprague-Dawley | $10^{5.07}$ TCID<sub>50</sub>/40 μL ($10^{6.47}$TCDI<sub>50</sub>/mL) |
| Wistar | $10^{5.18}$ TCID<sub>50</sub>/40 μL ($10^{6.58}$TCDI<sub>50</sub>/mL) |

TCID<sub>50</sub>: 50% Tissue Culture Infectious Dose

### Modelling of the interaction between the RBD of Omicron BA.5 and the ACE2

The percentage of identity (amino acids sequences) between angiotensin-converting enzyme 2 (ACE2) of both *Rattus* species was 96%. The *Homo sapiens* ACE2 shared respectively 82% and 79% of identity with *Rattus norvegicus* and *Rattus rattus* (S1 File). Positions of the Omicron BA.5 receptor binding domain (RBD) relative to ACE2 receptors of *Homo sapiens*, *Rattus norvegicus* and *Rattus rattus* are showed on Figs 7A-C. The affinity of each complex was estimated using PMF calculations based on molecular dynamics simulations, which required separating Omicron BA.5 from ACE2. Complete separation of both proteins were visualised by a plateau (Fig 7D), the resulting binding energy values for *Homo sapiens*, *Rattus norvegicus* and *Rattus rattus* were respectively 45 ± 1, 28 ± 1 and 33 ± 1 kcal/mol.

**Fig 7.**
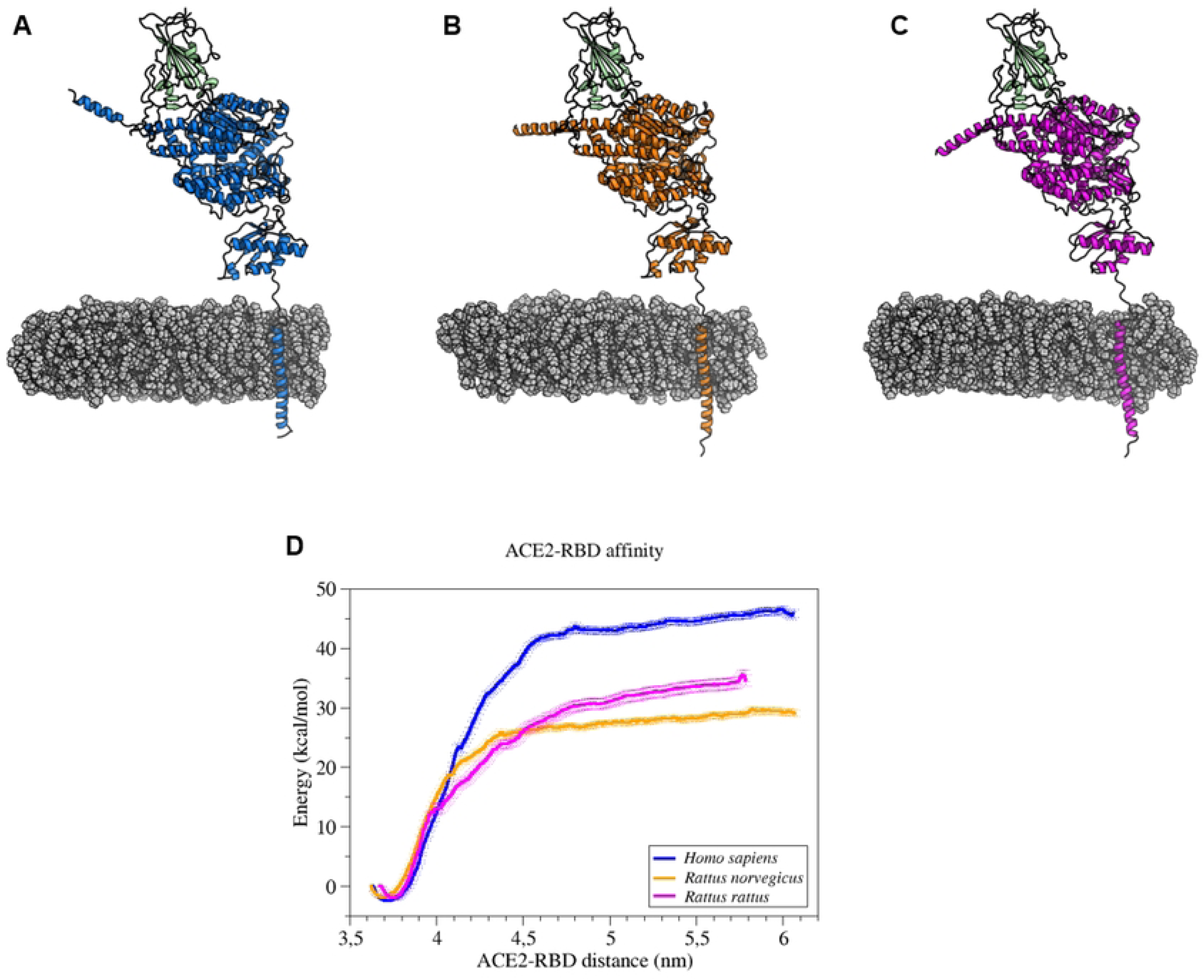
Modelling of the interaction between the receptor binding domain (RBD) of SARS-CoV-2 Omicron BA.5 and the ACE2 of *Homo sapiens*, *Rattus norvegicus* and *Rattus rattus*. 3-dimensions structures showing receptor cellular ACE2 of *Homo sapiens* (blue) (A), *Rattus norvegicus* (orange) (B) and *Rattus rattus* (pink) (C) interacting with the receptor binding domain (RBD) of Omicron BA.5 (green). Cell membrane is representing in grey. (D) ACE2/RBD affinities in kcal/mol during separation of both proteins with *Homo sapiens* in blue, *Rattus norvegicus* in orange and *Rattus rattus* in pink.

## Discussion

Our study primarily investigates the circulation of SARS-CoV-2 in wild rats, *Rattus norvegicus*, in France. Trapping was conducted in multiple cities throughout France, including Besançon, Nancy, Bordeaux, Lyon, Marseille, and Nantes, with the objective of enhancing the representativeness of the study population. Of the 401 rats included in the study, 351 were captured in areas connected to the sewer system, with 117 of these being captured directly within sewers. Sampling efforts were concentrated in sewers or areas in close proximity to sewers due to the extensive report of the SARS-CoV-2 genome in wastewater [23], which may serve as a potential source of infection for rats in close contact with contaminated water. The sampling was conducted between January 2022 and July 2023, a period during which several SARS-CoV-2 variants were circulating in human population (i.e. Omicron BA.2, BA.5, XBB 1.5 and EG.5). In addition, the effective presence of SARS-CoV-2 in the wastewater from the various trapping areas was confirmed to be present. The circulation of SARS-CoV-2 in rats was assessed using a dual approach that combined molecular biology techniques for detecting specific viral RNA in lungs, with serological analysis to detect specific anti-SARS-CoV-2 antibodies.

No traces of SARS-CoV-2 viral RNA were detected in any of the 401 lungs samples by RT-PCR, indicating the absence of viral replication in the lungs of the trapped rats. This finding is significant, as the lungs are the tissue where viral replication would typically be expected, and thus these results rule out the presence of an ongoing SARS-CoV-2 infection in these rats, despite the presence of the virus genome in the wastewater.

Similar results have been observed in previous studies, which also reported the absence of viral RNA in the lungs of wild rats captured in Belgium, Germany and Spain [24–26], as well as in other matrices, including feces, oronasopharyngeal swabs, nasal turbinates, intestines, and various other tissues [20,27,28]. Despite these results, infection of rats by SARS-CoV-2 seems possible, as a few rare studies have reported positive cases. For instance, four out of 79 rats tested positive in the United States, and three out of 72 in Mexico [15,29], but it should be noted that these results were obtained using RT-PCR methods different from the one carried out in our study (different targeted genes and primers). Several hypotheses can be put forward to explain our findings. Either SARS-CoV-2 was undetectable within rat populations in contact with wastewater during the study period. Or the virus present in the wastewater was not viable or infectious [30], which would prevent the rats from becoming contaminated, despite a recent study indicating that the virus can survive for a few hours in wastewater under experimental conditions [18]. Another possibility is that, although the rats were exposed to the virus, it was unable to replicate efficiently in their tissues due to mutations in the spike protein that could affect its affinity for the cell entry receptor or alter its replication in rodent, as previously shown for the SARS-CoV-2 B.1.351 variant [14], the B.1.1.529 Omicron (BA.1) [31,32] or the BA.5.5 Omicron variants [15].

To explore this further, serological analyses were conducted in parallel to detect potential signs of past infection. Of the 353 serum samples from *Rattus norvegicus*, three produced inconclusive results, testing positive for two out of three SARS-CoV-2 antigens when analyzed using a multiplex microsphere immunoassay. The three inconclusive samples later tested negative in the neutralization assay, confirming the absence of neutralizing SARS-CoV-2 antibodies in these rats. This may be explained by the absence of exposure of the trapped wild rats to infectious virus in wastewater, or by low seroconversion and the lack of maintenance of antibodies over time. Another possibility should be that the antibodies produced by the SARS-COV-2 variants circulating in wastewater at the time of the study are not able to neutralise the variant D614G used as challenge virus in our neutralisation assays. Indeed, as recent findings demonstrated that Omicron variants escape serum neutralisation to a greater extent than the ancestral D614G strain, we can put forward the opposite hypothesis that the D614G variant could be insensitive to antibodies raised against the new Omicron variants.

Our findings are in concordance with those of Colombo *et al*. obtained at the beginning of the pandemic, who reported that 3 out of 35 *Rattus norvegicus* were positive by MIA but were all negative by viral neutralisation test [24]. In addition, no SARS-CoV-2 antibodies were detected in studies conducted on wild rat populations in Germany, Belgium/France and Spain [25,26,33]. Conversely, the presence of antibodies by seroneutralisation assays has been reported in *Rattus sp.* in the USA (2 out of 117 *Rattus norvegicus*), Hong-Kong (1 out of 217 *Rattus norvegicus*), Canada (2 out of 213 *Rattus norvegicus*), Germany (1 out of 130 *Rattus norvegicus*) and Malaysia (1 out 3 *Rattus rattus*) [20,27,28,34,35]. However, some studies detected antibodies using other methods for detecting binding antibodies such as ELISA but reported negative results by neutralisation assays [15,19]. Seroneutralisation results can vary depending on the method used (with live virus or purified proteins) [36]. Indeed, studies demonstrating positive neutralising antibody samples surrogate virus neutralising test (sVNT) using proteins (mainly RBD) instead of live virus such as in our study. Furthermore, sera positive by MIA or ELISA were negative by conventional surrogate virus neutralising test (cVNT) using live virus [15,24]. cVNT is the gold standard to detect neutralising antibodies and a high correlation between these methods has been demonstrated [37].

Regardless of the method used, cross-reactivities of the detected SARS-CoV-2 antibodies with other coronaviruses circulating in rats cannot be excluded, as recently demonstrated for human and other animals [38]. This phenomenon was evoked for SARS-CoV-2 [15,20,28] and for coronaviruses detected in wild rodents [39].

In order to advance our understanding of the epidemiological role of wild rats – whether they are unexposed, unresponsive and or insensitive – we inoculated wild rats (*Rattus norvegicus* and *Rattus rattus*) and laboratory rats (Sprague-Dawley and Wistar) with the SARS-CoV-2 Omicron BA.5. This strain was the main SARS-CoV-2 variant that circulate during the sampling of wild rats.

No clinical signs or body weight loss were observed in the inoculated groups in comparison to the controls. These findings align with those of previous *in vivo* studies conducted on Sprague-Dawley rats infected with various SARS-CoV-2 variants, including the Wuhan-like strain, Alpha, Beta, Delta, and Omicron (BA.5) [12,14,15]. In our experiments, only viral RNA was isolated in the URT at 4 dpi, with RNA titers ranging between 4.7×10^3^ and 3.2×10^5^ RNA copies/mg of tissue, while no viral RNA was detected in the lungs, and no infectious particles (TCID50 assays) were detected in any tissue (lungs or URT). Our results in URT samples could be due to the difference of sensitivity between RT-qPCR and TCID50 assay (cell culture). Indeed, in our laboratory, the thresholds of detection are 10^2^ TCID50/mL and 25 copies/µL for the TCID50 assays and RT-qPCR tests respectively. It is also important to note that in our study, all animals were inoculated with a high dose of virus (titers ranging from 10^5.97^ to 10^6.74^ TCID50/mL (approximately 4.9×10^6^ to 5.5×10^6^ RNA copies/µL)), and viral RNA titers found in the URTs were systematically lower (between 3.3×10^2^ and 8.8×10^3^ RNA copies/µL of URT supernatant) than these virus doses. It could therefore also be assumed that only remnants of viral RNA or viral particles from the inoculum were detected by RT-qPCR at 4 dpi which could help to explain the absence of infectious virus in URT and lungs and viral RNA in lungs. These results provide further evidence to support the hypothesis that either viral replication occurred at a very low level in these tissues, or was abortive or absent. Discrepant results were observed in the existing literature by using a different Omicon variant. Indeed, for Omicron BA.5.5 variant, Wang *et al*. detected infectious virus only at 2 dpi in the lungs of Sprague-Dawley rats, inoculated with 2.10^4^ PFU/rats of virus, while no infectious particles were detected in URT [15]. In addition, in their study, viral RNA was detected at both 2 and 4 dpi in lungs and URT at a level demonstrating efficient replication [15].

Regarding the humoral immune response to the infection with Omicron BA.5 variant by intranasal (IN) route, it shall be noted that antibodies were detected in all animals of the two laboratory strains, whereas seroconversion was only observed in four of the eight wild *Rattus norvegicus* and four of the seven *Rattus rattus*. These differences between wild rats and laboratory rats can be attributed to the immune system specific to each of these models. In the case of wild rats, we need to talk about eco-immunology, because their immune response is conditioned by the environment and the external stimuli they are confronted with on a daily basis. As a reminder, the three key concepts of eco-immunology are immuno-heterogeneity, sub-maximal immune response due to energy limitation and the diversity of the antigenic load with which animals may be confronted. In the case of laboratory animals, they are bred under controlled conditions. Immunoheterogenicity is therefore limited, energy resources are not limited to maximise immune response and contact with external pathogens is kept to a minimum. Taken together, these factors could optimise their immune response, unlike wild animals [40]. The results obtained in the present study are in agreement with those reported by Wang et al., demonstrating the presence of IgG and neutralising antibodies at 21 dpi [15].

These results obtained for the *in vivo* manipulations, even for wild rats, are inconsistent with those obtained in the field. Indeed, contrary to the results obtained in our field study, the presence of anti-SARS-CoV-2 antibodies was confirmed in all four groups of rats at 14 dpi, using both MIA and serum neutralisation tests despite the absence of virus replication in the target tissues. A phenomenon of more or less marked passive immunisation of animals in response to the presence SARS-CoV-2 viral particles inoculated by IN route could explain the antibody production. This finding could also suggest that the trapped rats during our field study were not exposed to infectious particles of SARS-COV-2 in sewers as no antibodies were detected.

To summarise the results of our *in vivo* studies, no signs of viral infection were detected in wild-type or laboratory rats following inoculation of the Omicron BA.5 variant by the IN route and the antibody response should be considered more as a marker of passive immunisation response than of viral infection.

The results obtained from our field and *in vivo* studies suggest that rats have little or no receptivity or susceptibility to infection with SARS-CoV-2 (particularly with the Omicron BA.5 variant). As indicated earlier in the discussion, the low or inefficient replication of the virus demonstrated in this article could be explained by a low affinity between the rat ACE2 receptor (cell entry receptor) and the Omicron BA.5 variant. Indeed, it is well known that virus-receptor interaction is generally an effective determinant of virulence. To explore this hypothesis, molecular modelling has been conducted between the ACE2 of *Rattus norvegicus*, *Rattus rattus* and *Homo sapiens* and the Omicron BA.5 RBD. The findings align with the experimental outcomes, indicating a diminished affinity of the RBD for the ACE2 of both *Rattus* species (28 ± 1 and 33 ± 1 kcal/mol) in comparison to human ACE2 (45 ± 1 kcal/mol). These results offer a partial explanation for the lack of susceptibility of rats to Omicron BA.5. Indeed, a low affinity between the RBD receptor and ACE2 results in a weaker attachment of the virus to the cell. Nevertheless, a discrepancy exists between the results of *in silico* calculations and *in vivo* experiments, particularly with regard to the role of the animal immune system. In the context of the 3Rs, molecular modelling could be employed as a replacement for animal experimentation in the first instance. In this context, the evaluation of diverse RBDs from emerging SARS-CoV-2 variants in comparison with the ACE2 of *Rattus* sp. could serve as a pivotal approach to ascertain the potential for these variants to infect rats, obviating the necessity for animal experimentation.

The findings of this study indicate that wild or laboratory rats do not play an epidemiologic role in the SARS-CoV-2 pandemic, particularly in relation to the Omicron BA.5 variant. Indeed, no evidence of the presence of SARS-CoV-2 RNA was identified in the lungs and sera of rats captured in urban and sewage environments in several French cities. Furthermore, the results of the *in vivo* experiments and *in silico* calculations are in accordance with those of the field study. Indeed, the four groups of rats (wild *Rattus norvegicus*, wild *Rattus rattus*, and laboratory *Rattus norvegicus* Sprague-Dawley and Wistar) exhibited no susceptibility to infection by the SARS-CoV-2 Omicron BA.5. In light of the rapid evolution of the virus, its adaptation to new hosts and the emergence of SARS-CoV-2 variants, it is imperative to continue studying the circulation of SARS-CoV-2 in wild rodents in order to investigate the potential establishment of a wildlife reservoir. Furthermore, as a preliminary approach, molecular modelling could be conducted using emerging SARS-CoV-2 variants before proceeding with animal experimentation.

## Materials and Methods

### Ethics statements

The experimental protocols involving rats complied with the regulation 2010/63/CE of the European Parliament and of the council of 22 September 2010 on the protection of animals used for scientific purposes [41] and as transposed into French law [42].

*In vivo* experimental studies were approved by the Anses/ENVA/UPEC ethics committee and authorised by the French Ministry of Research (Apafis authorization n° #37782-2022062312057722 v4).

Field experimental studies were approved by the ethics committee of the Veterinary School of Lyon and authorised by the French Ministry of Research (Apafis authorization n° #37713-2022042712262465).

### Rat captures and samples

Wild rats were captured in sewers and parks connected or not to sewers in several French cities including Besançon, Bordeaux, Lyon, Marseille, Nancy and Nantes between January 2022 and July 2023. Chronologically, Omicron BA.2, BA.5, XBB 1.5 and EG.5 circulated in the French population during the sampling period [22]. *Rattus norvegicus* were captured using live traps or obtained through pest control campaigns. The rats were then transported in cages to a laboratory, where they were anaesthetised with isoflurane and euthanised by cardiac puncture and cervical dislocation. Various measurements including sex, weight, height, body length, and tail length were meticulously reported for each animal. Cytochrome b was performed to confirm the rat species in case of uncertainty. Each animal was aseptically autopsied and various tissues, including lung were removed, transferred to sterile tubes, and stored at −80°C until RNA extraction. For viral RNA extraction, lung tissue samples (∼20 to 60 mg) were placed in tubes containing beads (Lysing Matrix E, MP Biomedicals™) and 1 mL of Dulbecco’s Modified Eagle medium (DMEM) with 1% antibiotics (Penicillin/Streptomycin (P/S)). The samples were mixed using a bead beater (MP Biomedicals™ FastPrep-24™ 5G Bead Beating Grinder and Lysis System) at 4 m/s for 10 s, repeated three times with 90 s pauses on melting ice between each cycle. Homogenates were then clarified by centrifugation (2000 g, 10 min, 4°C). Supernatants were collected, aliquoted and stored at −80°C until analysis. Viral RNA was then extracted from 200 µL of supernatant using the Maxwell® Viral TNA kit, according to the manufacturer’s instructions (Promega, France). A negative control RNA extraction was performed for each set of 16 samples tested. All RNA extracts were stored at −80°C prior to use.

### Virus production

SARS-CoV-2 Omicron BA.5 was gracefully obtained from Pasteur Institut - National Reference Center for Viruses of Respiratory Infections (including Influenza and SARS-CoV-2) (virus sequence GISAID accession number EPI_ISL_13017789). The virus was passaged once on Calu-3 cells to increase the titer and the quantity available for animal infection studies. The NGS virus sequence obtained after passage on Calu-3 cells was 100% similar to the sequence of the virus stock.

### Experimental design

Fifty-two wild rats with uncontrolled genetic background and microbiota were included in this experimental study. This comprised twenty wild *Rattus rattus* (black rats) obtained from a colony that has been maintained in an outdoor enclosure for over 20 years at the veterinary school of Lyon (France). In the following, this group will be referred to as ‘*Rattus rattus*’. Additionally, thirty-two wild *Rattus norvegicus* (brown rats) were randomly selected from three distinct colonies A, B and C, each maintained in outdoor enclosures for over 20 years at the same facility. Animals from three different colonies were used in order to increase genetic diversity (the founder animals having been taken from different farms in the Lyon region). In the following, this group will be referred to as ‘wild *Rattus norvegicus*’. In addition to these wild specimens, forty laboratory rats with a known genetic background, consisting of twenty Sprague-Dawley strain and twenty Wistar strain, were included in the study. In the following, these groups will be referred to as ‘Sprague-Dawley’ and ‘Wistar’. Rats were kept in cages with environmental enrichment and housed either individually (wild *Rattus norvegicus*) or in groups of two rats per cage. Negative animals were kept in separate room from the SARS-CoV-2 inoculated animals. Food and water were provided *ad libitum*. Weight of all animals were monitored and recorded on a daily basis throughout the duration of the experimental procedures.

### Virus inoculation

Twenty-three wild *Rattus norvegicus*, fourteen wild *Rattus rattus*, fourteen Sprague-Dawley and fourteen Wistar were anaesthetized with isoflurane and intranasally inoculated (20µl in each nostril) with viral titters from 10^4.57^ TCID50 (50% Tissue Culture Infectious Dose)/40 µL to 10^5.34^ TCID50/40µL of SARS-CoV-2 Omicron BA.5 (Table 3). Nine wild *Rattus norvegicus*, six wild *Rattus rattus*, six Sprague-Dawley and six Wistar were inoculated with PBS in parallel to act as negative controls for each experimental protocol. As described on Fig 8, SARS-CoV-2 and PBS inoculated rats were euthanised at either 4 or 14 days post-inoculation (dpi).

**Fig 8.**
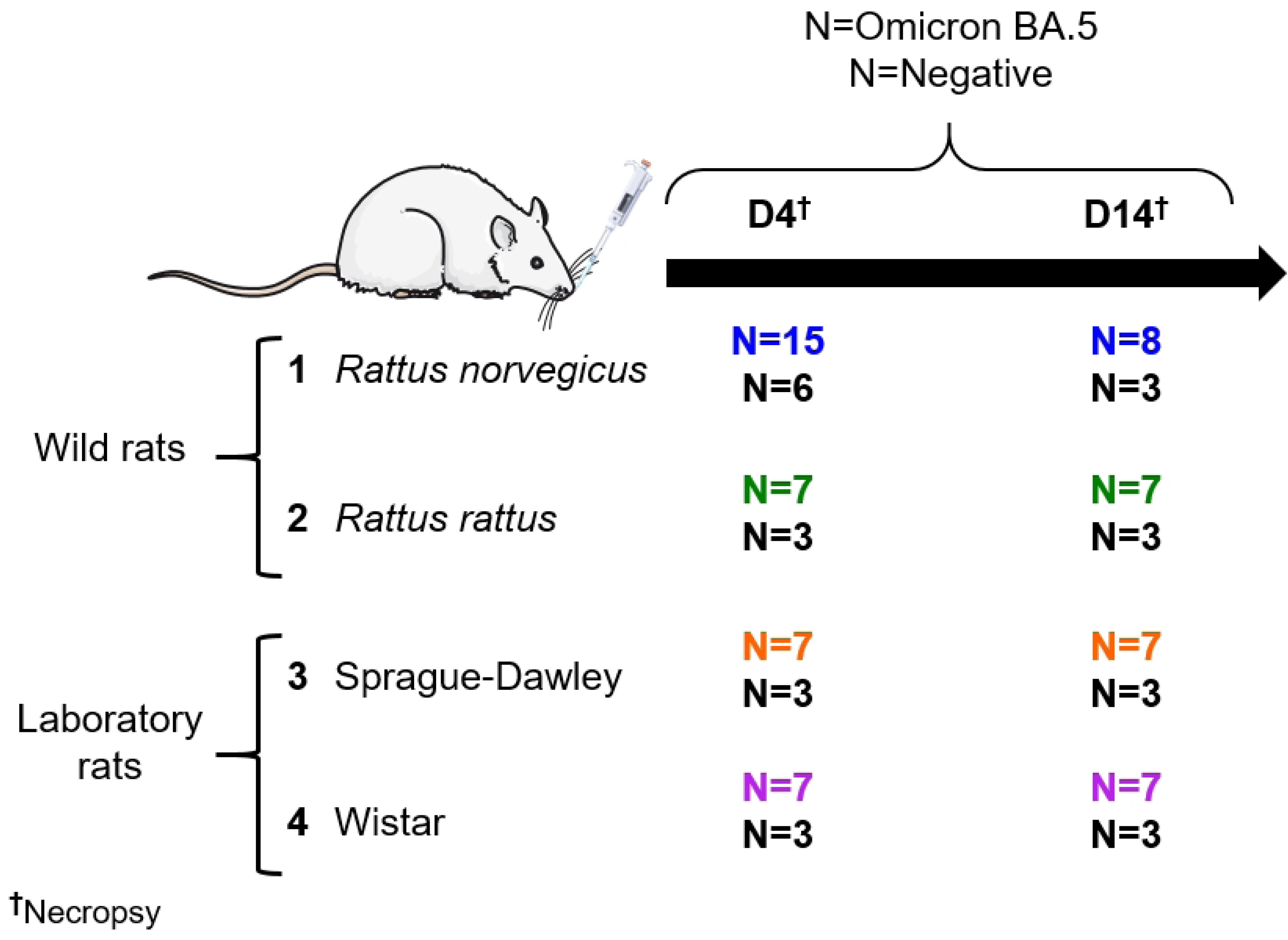
Experimental design including four groups of rats intranasally inoculated with Omicron BA.5. Wild *Rattus norvegicus* (blue), wild *Rattus rattus* (green), Sprague-Dawley laboratory rats (orange) and Wistar laboratory rats (purple), each inoculated with either SARS-CoV-2 Omicron BA.5 or PBS (negative control groups (black)) were necropsied at 4 (D4) or 14 (D14) days post infection.

### *In vivo* experiments and samples

Blood was collected by cardiac puncture under deep anaesthesia (Isoflurane, Ketamine (100 mg/kg) and Xylazine (10 mg/kg)) prior to euthanasia. Blood was collected into 4 mL K3-EDTA tubes. Following euthanasia, all animals were necropsied to collect lungs and upper respiratory tract (URT). After centrifugation of blood samples (1000 g/15 min), plasmas were stored at −20°C until analysis. For viral RNA extraction, lungs or URT tissues (∼20 to 60 mg) were prepared as described previously (section rat captures and samples). Viral RNA was extracted from 160 µL of previously obtained homogenate using the Qiagen Viral RNA mini kit (Qiagen, Courtaboeuf, France) according to the manufacturer’s instructions. All RNA extracts were stored at −80°C before use.

### SARS-CoV-2 viral titrations on cell culture

SARS-CoV-2 titrations were performed on lung and URT samples by TCID50 assays on Vero E6 cells.

Briefly, 96-well plates were seeded with a Vero E6 cell suspension (1.10^5^ cells/mL) 24 h before virus inoculation. A ten-fold serial dilution of tissue homogenates and the reference virus (used as positive control) was performed in DMEM supplemented with 10% Foetal Calf Serum (FCS) and 1% antibiotics (e.g. mix of P/S). After removing the cell culture medium, inoculum was added to each well and plates were incubated at 37°C in 5% CO_2_ for 1 h. Six uninfected wells were used as independent negative controls. After the 1 h incubation, 200 µL of DMEM supplemented with 10% FCS and 1% mix of P/S were added to each well and the plates were incubated at 37°C in 5% CO_2_ for 72 h. Viral titers were calculated in TCID50/mL using the Spearman-Kärber method.

In addition, viral inoculums were titrated on cells both before and after rat inoculations.

### Microsphere immunoassay (Luminex Technology)

For plasma samples from experimental *in vivo* studies and sera from field studies, anti-SARS-CoV-2 antibodies were detected by microsphere immunoassay (MIA) as described by Bourret *et al*. [33]. Three recombinant SARS-CoV-2 antigens (nucleoprotein, spike subunit 1 receptor binding domain (RBD) and trimeric spike (its native form) (The Native Antigen Company)) were used to capture specific plasma/serum antibodies. BSA (Bovine Serum Albumin) was used as a control antigen. Plasma samples from *in vivo* studies were previously diluted 1:20 while sera from field studies were diluted 1:10 in assay buffer (PBS-1% BSA-0.05% Tween 20, (Sigma-Aldrich)). The assay was performed on a Bio-Plex 200 instrument (Bio-Rad). A minimum of 100 events were read for each bead set and binding events were expressed as median fluorescence intensity (MFI). Relative Fluorescence Intensity (RFI) was calculated for each sample by dividing the MFI signal measured for the antigen coated microsphere sets by the MFI signal obtained for the control microsphere set (BSA coated beads), to account for non-specific binding of antibodies to the beads. Specific seropositivity cut-off values for each antigen were set at three standard deviations above the mean RFI of negative samples. To determine the seropositivity cut-off for the field samples, given the low prevalence of SARS-CoV-2 in wild rodents, the entire rat population (mean of all RFIs + 3 standard deviation) was used as described by Fritz *et al.* [21]. Seroconversion was defined as the detection of antibodies to the three antigens and inconclusive result was defined as the detection of antibodies to two of the three antigens.

### Seroneutralisation assay

Seroneutralisation assays were carried out on heat-inactivated plasmas/sera (56°C/30 min) collected from all rats included in the *in vivo* experiments, as well as on positive/inconclusive sera by MIA collected from urban rodents. The SARS-CoV-2 Omicron BA.5 (virus sequence GISAID accession number EPI_ISL_13017789) and the SARS-CoV-2 D614G (virus sequence GISAID accession number EPI_ISL_666870) variants were used respectively as challenge viruses as described by Bourret *et al*. [33]. Briefly, Vero E6 cells were seeded in 96-well plates at 1.10^5^ cells/mL. One day later, plasmas, positive and negative controls were serially diluted in a 1 to 3 dilution steps in cell culture medium (DMEM supplemented with 10% SVF and 1% P/S) and 50 µL of virus previously diluted to 100 TCID50/50µL were added to samples and controls in the 96-well plates. The plates were incubated at 37°C (5% CO_2_) for 1 h. Afterwards, the cell culture supernatants were removed and replaced with 100 µL of the mix of virus and serially diluted samples or controls. After at least 3 days of incubation, the presence of viral cytopathic effect (CPE) was quantified by an “all or nothing” reading method. The neutralisation titer was calculated from the highest dilution that prevented detectable cytopathic effect.

### TaqMan RT-qPCR

TaqMan RT-qPCR targeting the envelope protein gene (E gene) was performed using the Quantitect Probe RT-PCR kit (Qiagen, Courtaboeuf, France). The model of primers/probe used was as follows: primer E_Sarbeco_F (forward): 5’-ACAGGTACGTTAATAGTTAATAGCGT, primer E_Sarbeco_R (reverse): 5’-ATATTGCAGCAGTACGCACACA and the probe E_Sarbeco_P1 5’-FAM-ACACTAGCCATCCTTA CTGCGCTTCG-BHQ-1. Primers and probe were supplied by Eurogentec (Angers, France).

PCR was performed as previously described in Monchatre Leroy *et al*. with minor modifications [43]. TaqMan RT-qPCR was performed in a total volume of 20 µL containing 2.5 μL of RNA sample, 10 µL of 2x Master Mix, 5.3 µL of RNase-free water, 0.8 µL each of forward and reverse primer (10 µM), 0.4 µL of probe (10 µM), and 0.2 μL of QuantiTect RT Mix. All TaqMan RT-qPCR assays were performed on the thermocycler Rotor Gene Q MDx (Qiagen, Courtaboeuf, France). Amplification was performed under the following thermocycling conditions: 50°C for 30 min for reverse transcription, followed by 95°C for 15 min and then 45 cycles of 94°C for 15 s and 60°C for 60 s. Negative and positive controls were included in each RT-qPCR assay.

Copy number/µL in each sample was determined using si× 10-fold serial dilutions of SARS COV-2 RNA at 3.23×10^8^ copies/µL (3.23×10^7^ - 3.23×10^2^). A cut-off of 0.03 was used as the reference threshold for each RT-qPCR assay. Efficiency, slope and correlation coefficient (R^2^) were calculated directly by the Rotor Gene software. All reactions were performed in duplicates.

Finally, the PCR detection limit was determined by testing five serial dilutions of SARS-COV-2 from 500 to 5 copies/µL, each dilution being performed in quadruplicate (i.e. 12 replicates per dilution). LD_PCR_ was demonstrated > 25 copies/µL (last dilution giving 100% positive replicates).

SARS-CoV-2 RT-qPCR, targeting the polymerase gene, was performed on purified nucleic acids extracted from sewage collected in sewers as described in [44] for Nancy, Besançon, Nantes and Lyon where rodent were trapped directly in sewers.

### Molecular modelling procedures

The complete structures (including the transmembrane domain) of *Homo sapiens* and *Rattus norvegicus* ACE2 receptors were obtained from AlphaFold protein structure database [45,46] and can be found using their UniProt identifiers (Q9BYF1 and Q5EGZ1, respectively). The structural model of the *Rattus rattus* ACE2 receptor was obtained using AlphaFold Colab notebook [47] based on the GenBank sequence (XM_032890254.1). The Receptor Binding Domain (RBD) structure of the Omicron BA.5 Spike protein was obtained from the Protein Data Bank [48], PDB ID: 7XWA [49]. ACE2-RBD complexes were finally built by superimposing their respective structures on the crystallographic structure of the above complex (7XWA), using PyMOL software (PyMOL).

Molecular dynamics (MD) systems preparation and simulations were performed using the GROMACS 2018 package software [50], with the CHARMM-36 forcefield applied [51]. Each ACE2-RBD complex was first embedded into a previously equilibrated phosphatidylcholine (DLPC) membrane [52] using the *membed* function of the GROMACS software. Solvation, energy minimization, 100-ps temperature and 1-ns pressure equilibrations were carried out using GROMACS routines (V-rescale thermostat, Berendsen pressure coupling, and semi-isotropic coupling type were applied). A 500-ns MD simulation was performed on each ACE2-RBD complex to equilibrate the systems. The *cluster* function of GROMACS, with the *gromos* algorithm and a 4 Å cut-off, was then applied to identify the most representative conformation of each complex, finally used for Potential of Mean Force (PMF) calculations.

PMF calculations involved ACE2 and RBD as pull groups (positions of the ACE2 receptors were restrained), which defined the Z-axis as the reaction coordinate. A harmonic potential and a distance increase over this reaction coordinate were then applied, with a constant velocity of 100 Å/ns and a force constant of 1000 kJ/mol/nm². After this pull step, the RBD positions along the reaction coordinate were sampled every 2 Å, until the RBD center of mass reached a 60 Å distance from the ACE2 receptor centre of mass. The Umbrella Sampling (US) step was then performed on the 11 resulting conformations, with a 1000 kJ/nm/nm² constant force over 4-ns simulations. Finally, the PMF values were extracted using the weighted histogram analysis method implemented in GROMACS. Percentages of identity between ACE2 sequences were calculated using Geneious Prime® 2022.0.2 version.

### Statistical analyses and graphs

Graphs and statistical analyses (Kruskall-Wallis and one-way ANOVA) were performed with GraphPad Prism (version 9) and PyMOL (version 4.6.0). A p-value of <0.05 was considered statistically significant. Figs 1 and 8 were generated using images from Servier Medical Art by Servier, licensed under a Creative Commons Attribution 4.0 Unported License (https://creativecommons.org/licenses/by/4.0/).

## Acknowledgements

We would like to thanks all the partners from urban cities involved in the pest control programs, the teams of ANSES Laboratory for Rabies and Wildlife including Virologie Moléculaire (C. Peytavin De Garam, F. Bastien and J.L. Schereffer) and Virologie-Immunologie-Sérologie (J. Rieder, A. Labadie, F. Chanteclair and J. Bonetti) and Service d’Expérimentation Animale Rongeurs (S. Kempff, V. Brogat, and E. Litaize) and Service d’Expérimentation Animale Carnivores (M. Vesvres). We would like to thanks Sylvie Van der Werf and Flora Donati from Pasteur Institut - National Reference Center for Viruses of Respiratory Infections (including Influenza and SARS-CoV-2) for providing SARS-CoV-2 Omicron BA.5.

## Funding

This work was funded by l’ANRS/Santé Publique France - Projet EMERGEN (ANRS0163). This work was granted access to the HPC resources of IDRIS under the allocation AD010715471 made by GENCI.

## Supporting information

**S1 Fig. Alignments of *Homo sapiens, Rattus norvegicus and Rattus rattus* ACE2.**

## Notes

### Competing Interest Statement

The authors have declared no competing interest.

## References

1. Zhou P, Yang XL, Wang XG, Hu B, Zhang L, Zhang W, et al. A pneumonia outbreak associated with a new coronavirus of probable bat origin. Nature. 2020;579(7798):270–3.

2. Qiu X, Liu Y, Sha A. SARS-CoV-2 and natural infection in animals. J Med Virol. 2022 Sep 28;10.1002/jmv.28147.

3. Sailleau C, Dumarest M, Vanhomwegen J, Delaplace M, Caro V, Kwasiborski A, et al. First detection and genome sequencing of SARS-CoV-2 in an infected cat in France. Transbound Emerg Dis. 2020 Jun 5;10.1111/tbed.13659.

4. Sit TH, Brackman CJ, Ip SM, Tam KW, Law PY, To EM, et al. Canine SARS-CoV-2 infection. Nature. 2020 Oct;586(7831):776–8.

5. Oreshkova N, Molenaar RJ, Vreman S, Harders F, Oude Munnink BB, Hakze-van der Honing RW, et al. SARS-CoV-2 infection in farmed minks, the Netherlands, April and May 2020. Euro Surveill Bull Eur Sur Mal Transm Eur Commun Dis Bull. 2020 Jun;25(23):2001005.

6. McAloose D, Laverack M, Wang L, Killian ML, Caserta LC, Yuan F, et al. From People to Panthera: Natural SARS-CoV-2 Infection in Tigers and Lions at the Bronx Zoo. mBio. 2020 Oct 13;11(5):e02220–20.

7. Padilla-Blanco M, Aguiló-Gisbert J, Rubio V, Lizana V, Chillida-Martínez E, Cardells J, et al. The Finding of the Severe Acute Respiratory Syndrome Coronavirus (SARS-CoV-2) in a Wild Eurasian River Otter (Lutra lutra) Highlights the Need for Viral Surveillance in Wild Mustelids. Front Vet Sci. 2022 Mar 31;9:826991.

8. Kuchipudi SV, Surendran-Nair M, Ruden RM, Yon M, Nissly RH, Vandegrift KJ, et al. Multiple spillovers from humans and onward transmission of SARS-CoV-2 in white-tailed deer. Proc Natl Acad Sci U S A. 2022 Feb 8;119(6):e2121644119.

9. Larsen HD, Fonager J, Lomholt FK, Dalby T, Benedetti G, Kristensen B, et al. Preliminary report of an outbreak of SARS-CoV-2 in mink and mink farmers associated with community spread, Denmark, June to November 2020. Euro Surveill Bull Eur Sur Mal Transm Eur Commun Dis Bull. 2021 Feb;26(5):2100009.

10. Pickering B, Lung O, Maguire F, Kruczkiewicz P, Kotwa JD, Buchanan T, et al. Divergent SARS-CoV-2 variant emerges in white-tailed deer with deer-to-human transmission. Nat Microbiol. 2022 Dec;7(12):2011–24.

11. Bao L, Deng W, Huang B, Gao H, Liu J, Ren L, et al. The pathogenicity of SARS-CoV-2 in hACE2 transgenic mice. Nature. 2020 Jul;583(7818):830–3.

12. Shuai H, Chan JFW, Yuen TTT, Yoon C, Hu JC, Wen L, et al. Emerging SARS-CoV-2 variants expand species tropism to murines. EBioMedicine. 2021 Nov;73:103643.

13. Muñoz-Fontela C, Dowling WE, Funnell SGP, Gsell PS, Balta XR, Albrecht RA, et al. Animal models for COVID-19. Nature. 2020 Oct;586(7830):509–15.

14. Zhang C, Cui H, Li E, Guo Z, Wang T, Yan F, et al. The SARS-CoV-2 B.1.351 Variant Can Transmit in Rats But Not in Mice. Front Immunol. 2022;13:869809.

15. Wang Y, Lenoch J, Kohler D, DeLiberto TJ, Tang CY, Li T, et al. SARS-CoV-2 Exposure in Norway Rats (Rattus norvegicus) from New York City. mBio. 2023 Apr 25;14(2):e0362122.

16. Montagutelli X, Prot M, Levillayer L, Salazar EB, Jouvion G, Conquet L, et al. Variants with the N501Y mutation extend SARS-CoV-2 host range to mice, with contact transmission. BioRxiv [Preprint]. 2021; 2021.03.18.436013 [posted 2021 Dec 07; cited 2024 Dec 10]: [22 p]. Available from: https://www.biorxiv.org/content/10.1101/2021.03.18.436013v2 doi: 10.1101/2021.03.18.436013

17. Smyth DS, Trujillo M, Gregory DA, Cheung K, Gao A, Graham M, et al. Tracking cryptic SARS-CoV-2 lineages detected in NYC wastewater. Nat Commun. 2022 Feb 3;13(1):635.

18. Sherchan S, Thakali O, Ikner LA, Gerba CP. Survival of SARS-CoV-2 in wastewater. Sci Total Environ. 2023 Jul 15;882:163049.

19. Fisher AM, Airey G, Liu Y, Gemmell M, Thomas J, Bentley EG, et al. The ecology of viruses in urban rodents with a focus on SARS-CoV-2. Emerg Microbes Infect. 2023 Dec 31;12(1):2217940.

20. Robinson SJ, Kotwa JD, Jeeves SP, Himsworth CG, Pearl DL, Weese JS, et al. Surveillance for SARS-CoV-2 in Norway Rats (Rattus norvegicus) from Southern Ontario. Wang L, editor. Transbound Emerg Dis. 2023 May 26;2023:1–9.

21. Fritz M, de Riols de Fonclare D, Garcia D, Beurlet S, Becquart P, Rosolen SG, et al. First Evidence of Natural SARS-CoV-2 Infection in Domestic Rabbits. Vet Sci. 2022 Jan 27;9(2):49.

22. Coronavirus : circulation des variants du SARS-CoV-2 [Internet]. [cited 2024 Dec 10]. Available from: https://www.santepubliquefrance.fr/dossiers/coronavirus-covid-19/coronavirus-circulation-des-variants-du-sars-cov-2

23. Wurtzer S, Waldman P, Levert M, Cluzel N, Almayrac JL, Charpentier C, et al. SARS-CoV-2 genome quantification in wastewaters at regional and city scale allows precise monitoring of the whole outbreaks dynamics and variants spreading in the population. Sci Total Environ. 2022 Mar 1;810:152213.

24. Colombo VC, Sluydts V, Mariën J, Vanden Broecke B, Van Houtte N, Leirs W, et al. SARS-CoV-2 surveillance in Norway rats (Rattus norvegicus) from Antwerp sewer system, Belgium. Transbound Emerg Dis. 2021 Jul 10;10.1111/tbed.14219.

25. Wernike K, Drewes S, Mehl C, Hesse C, Imholt C, Jacob J, et al. No Evidence for the Presence of SARS-CoV-2 in Bank Voles and Other Rodents in Germany, 2020–2022. Pathogens. 2022 Sep 28;11(10):1112.

26. Fernández-Bastit L, Montalvo T, Franco S, Barahona L, López-Bejar M, Carbajal A, et al. Monitoring SARS-CoV-2 infection in urban and peri-urban wildlife species from Catalonia (Spain). One Health Outlook. 2024 Sep 1;6(1):15.

27. Lee LKF, Himsworth CG, Prystajecky N, Dibernardo A, Lindsay LR, Albers TM, et al. SARS-CoV-2 Surveillance of Wild Mice and Rats in North American Cities. EcoHealth. 2024 Mar;21(1):1–8.

28. Miot EF, Worthington BM, Ng KH, Lataillade L de G de, Pierce MP, Liao Y, et al. Surveillance of Rodent Pests for SARS-CoV-2 and Other Coronaviruses, Hong Kong. Emerg Infect Dis J - CDC. 2022 Feb 1;28(2):467–470.

29. Martínez-Hernández F, Gonzalez-Arenas NR, Cervantes JAO, Villalobos G, Olivo-Diaz A, Rendon-Franco E, et al. Identification of SARS-CoV-2 in urban rodents from Southern Mexico City at the beginning of the COVID-19 pandemic. Rev Inst Med Trop São Paulo. 2024 Feb 5;66:e8.

30. Robinson CA, Hsieh HY, Hsu SY, Wang Y, Salcedo BT, Belenchia A, et al. Defining biological and biophysical properties of SARS-CoV-2 genetic material in wastewater. Sci Total Environ. 2022 Feb 10;807(Pt 1):150786.

31. Halfmann PJ, Iida S, Iwatsuki-Horimoto K, Maemura T, Kiso M, Scheaffer SM, et al. SARS-CoV-2 Omicron virus causes attenuated disease in mice and hamsters. Nature. 2022;603(7902):687–92.

32. Shuai H, Chan JFW, Hu B, Chai Y, Yuen TTT, Yin F, et al. Attenuated replication and pathogenicity of SARS-CoV-2 B.1.1.529 Omicron. Nature. 2022 Mar;603(7902):693–9.

33. Bourret V, Dutra L, Alburkat H, Mäki S, Lintunen E, Wasniewski M, et al. Serologic Surveillance for SARS-CoV-2 Infection among Wild Rodents, Europe. Emerg Infect Dis J. 2022;28(12):2577–80.

34. Wernike K, Mehl C, Aebischer A, Ulrich L, Heising M, Ulrich RG, et al. SARS-CoV-2 and Other Coronaviruses in Rats, Berlin, Germany, 2023. Emerg Infect Dis. 2024 Oct;30(10):2205–8.

35. Tan CS, Adrus M, Rahman SPH, Azman HIM, Abang RAA. Seroevidence of SARS-CoV-2 spillback to rodents in Sarawak, Malaysian Borneo. BMC Vet Res. 2024 Apr 27;20(1):161.

36. Perera RAPM, Ko R, Tsang OTY, Hui DSC, Kwan MYM, Brackman CJ, et al. Evaluation of a SARS-CoV-2 Surrogate Virus Neutralization Test for Detection of Antibody in Human, Canine, Cat, and Hamster Sera. J Clin Microbiol. 2021 Jan 21;59(2):e02504–20.

37. Mariën J, Michiels J, Heyndrickx L, Nkuba-Ndaye A, Ceulemans A, Bartholomeeusen K, et al. Evaluation of a surrogate virus neutralization test for high-throughput serosurveillance of SARS-CoV-2. J Virol Methods. 2021 Nov;297:114228.

38. Hulst M, Kant A, Harders-Westerveen J, Hoffmann M, Xie Y, Laheij C, et al. Cross-Reactivity of Human, Wild Boar, and Farm Animal Sera from Pre- and Post-Pandemic Periods with Alpha- and Βeta-Coronaviruses (CoV), including SARS-CoV-2. Viruses. 2023 Dec 23;16(1):34.

39. Lau SKP, Woo PCY, Li KSM, Tsang AKL, Fan RYY, Luk HKH, et al. Discovery of a Novel Coronavirus, China Rattus Coronavirus HKU24, from Norway Rats Supports the Murine Origin of Betacoronavirus 1 and Has Implications for the Ancestor of Betacoronavirus Lineage A. J Virol. 2014 Dec 31;89(6):3076–92.

40. Viney M, Riley EM. The Immunology of Wild Rodents: Current Status and Future Prospects. Front Immunol. 2017;8:1481.

41. Directive 2010/63/EU of the European Parliament and of the Council of 22 September 2010 on the protection of animals used for scientific purposes Text with EEA relevance [Internet]. 2010 [cited 2024 Dec 10]. Available from: http://data.europa.eu/eli/dir/2010/63/oj/eng

42. Article 1 - Décret n° 2013-118 du 1er février 2013 relatif à la protection des animaux utilisés à des fins scientifiques - Légifrance [Internet]. [cited 2024 Dec 10]. Available from: https://www.legifrance.gouv.fr/jorf/article_jo/JORFARTI000027037854

43. Monchatre-Leroy E, Lesellier S, Wasniewski M, Picard-Meyer E, Richomme C, Boué F, et al. Hamster and ferret experimental infection with intranasal low dose of a single strain of SARS-CoV-2. J Gen Virol. 2021 Mar;102(3):001567.

44. Vincent-Hubert F, Wacrenier C, Desdouits M, Jousse S, Schaeffer J, Le Mehaute P, et al. Development of passive samplers for the detection of SARS-CoV-2 in sewage and seawater: Application for the monitoring of sewage. Sci Total Environ. 2022 Aug 10;833:155139.

45. Jumper J, Evans R, Pritzel A, Green T, Figurnov M, Ronneberger O, et al. Highly accurate protein structure prediction with AlphaFold. Nature. 2021 Aug;596(7873):583–9.

46. Varadi M, Bertoni D, Magana P, Paramval U, Pidruchna I, Radhakrishnan M, et al. AlphaFold Protein Structure Database in 2024: providing structure coverage for over 214 million protein sequences. Nucleic Acids Res. 2024 Jan 5;52(D1):D368–75.

47. Mirdita M, Steinegger M, Söding J. MMseqs2 desktop and local web server app for fast, interactive sequence searches. Bioinformatics. 2019 Aug 15;35(16):2856–8.

48. Berman HM, Westbrook J, Feng Z, Gilliland G, Bhat TN, Weissig H, et al. The Protein Data Bank. Nucleic Acids Res. 2000 Jan 1;28(1):235–42.

49. Kimura I, Yamasoba D, Tamura T, Nao N, Suzuki T, Oda Y, et al. Virological characteristics of the SARS-CoV-2 Omicron BA.2 subvariants, including BA.4 and BA.5. Cell. 2022 Oct;185(21):3992–4007.e16.

50. Lemkul JA. From Proteins to Perturbed Hamiltonians: A Suite of Tutorials for the GROMACS-2018 Molecular Simulation Package [Article v1.0]. Living J Comput Mol Sci. 2019;1(1):5068–5068.

51. Best RB, Zhu X, Shim J, Lopes PEM, Mittal J, Feig M, et al. Optimization of the additive CHARMM all-atom protein force field targeting improved sampling of the backbone φ, ψ and side-chain χ(1) and χ(2) dihedral angles. J Chem Theory Comput. 2012 Sep 11;8(9):3257–73.

52. Jämbeck JPM, Lyubartsev AP. Derivation and systematic validation of a refined all-atom force field for phosphatidylcholine lipids. J Phys Chem B. 2012 Mar 15;116(10):3164–79.

